# An oscillator ensemble model of sequence learning

**DOI:** 10.1101/645085

**Authors:** Alexander Maye, Peng Wang, Jonathan Daume, Xiaolin Hu, Andreas K. Engel

## Abstract

Learning and memorizing sequences of events is an important function of the human brain and the basis for forming expectations and making predictions. Learning is facilitated by repeating a sequence several times, causing rhythmic appearance of the individual sequence elements. This observation invites to consider the resulting multitude of rhythms as a spectral ‘fingerprint’ which characterizes the respective sequence. Here we explore the implications of this perspective by developing a neurobiologically plausible computational model which captures this ‘fingerprint’ by attuning an ensemble of neural oscillators. In our model, this attuning process is based on a number of oscillatory phenomena that have been observed in electrophysiological recordings of brain activity like synchronization, phase locking and reset as well as cross-frequency coupling. We compare the learning properties of the model with behavioral results from a study in human participants and observe good agreement of the errors for different levels of complexity of the sequence to be memorized. Finally, we suggest an extension of the model for processing sequences that extend over several sensory modalities.

## 1 Introduction

Oscillations are a ubiquitous phenomenon when brain activity is observed at a sufficiently high temporal resolution, e.g., using EEG/MEG (electro-/magneto-encephalography) or invasive methods. Great progress has been made towards understanding the functional role of oscillations in cognitive processes [4,6–8,11, 20]. Their rhythmic nature suggests that neuronal oscillations could be used by the brain for learning, recognizing and producing rhythmic patterns in the interaction with the environment, and corresponding mechanisms have been suggested and studied in computational models. In particular, oscillator-based models have replicated many of the properties of human memory for serial order [3].

Probably the two most influential computational mechanisms are the encoding of arbitrary time intervals by an ensemble of oscillators with different periods and the dynamic adjustment of oscillation frequency and phase. The time representation by a single oscillator is limited by its period length and phase resolution. In a set of oscillators with different frequencies and phases however more rapid oscillations can provide temporal accuracy, while slower oscillations disambiguate cycles of the faster oscillations [5]. Basically the phases of the oscillators in the set provide a unique temporal context which can be associated with a sequence of events in the environment [3]. This dynamic context has a number of desirable properties for learning sequences of events: First, despite the cyclic activity of the individual oscillators, the vector of the combined phases is non-repetitive or at least repeats over very long epochs. By associating items in a complex sequence (e.g., ABAC) with the dynamic learning context, repetitions of the same item can be disambiguated. Second, the learning context for adjacent time points, when only the phases of oscillators with higher frequencies made substantial progress, is more similar than between more distant points, when also the phases of the low-frequency rhythms progressed. This property makes the approach suitable for sequences that involve temporal hierarchies like, for example, spoken language. And third, the series of learning contexts can easily be replayed by resetting the oscillators to their initial phase and restarting the clocking. By modifying the scale of the time signal that drives the oscillators in the set, stored sequences can be replayed at rates that are different from the original one.

The dynamic adjustment of oscillation frequency and phase is another mechanism which is frequently employed in computational models. The main idea is that the phase of the input relative to the ongoing oscillations determines how the synchronization patterns between the neural populations change. Sudden changes of the phase of ongoing oscillations in response to a stimulation, so called phase resetting, can frequently be observed in signals from human and animal brains, where it is considered to underlie multisensory integration functions [16, 21]. The simultaneous tuning of phase and frequency is aptly modeled by a phase-locked loop (PLL), in which the phase difference between an external rhythm and the ongoing oscillation generates a signal that adjusts the PLL’s frequency to minimize this phase difference. In PLL-based computational models of neuronal processing, memorized patterns are not equilibria or attractor states, like in conventional artificial neural networks, but synchronized oscillatory states with a certain phase relation [13]. The dynamically stable oscillation patterns can dynamically bind and unbind neural populations by synchronization, which can be used to model cognitive processes in working memory for associating and dissociating elements, inference by binding objects to the variables of a predicate, or algebraic operations defined by the transition rules between oscillation patterns of the network [18].

In this article we introduce a new perspective on sequence learning and present a computational model which integrates the two mechanisms of information processing by oscillatory dynamics that were discussed above. This perspective rests on the observation that when humans learn sequences, they frequently do so by verbally or mentally repeating the sequence over and over again. For example, to memorize the number code 99392, one might repeat ‘99392 99392 99392…’ a few times, e.g., by reading it off again from a note or mentally rehearsing it from short-term memory. This repetition can entrain a rhythm for each item. In the example, appearances of the digit ‘9’ would entrain a high frequency rhythm, whereas the rhythms entrained by digits ‘3’ and ‘2’ would have have lower frequencies and distinct phases. In addition to the periods that correspond to the temporal distance between any two repeating items, even slower rhythms can emerge when items in every other repetition are considered, whereas fast rhythms could cycle several times between two successive appearances of an item. All the different rhythms that are entrained by this sequence together constitute a characteristic entity that can be used to recognize correct instantiations of the sequence and detect deviations. Any incongruent item, e.g., the erroneous ‘2’ at the end of ‘99392 92’, would disturb the rhythms that were entrained by digits ‘2’ and ‘9’ during the learning phase and would be easily detected. From this perspective, the rhythms of a sequence appear to be analogous to the polyphony of an orchestra in which the tempi of the individual instruments compose an integrated experience that is unique for the respective piece of music and that is easily impaired by one or several instruments getting out of tune.

In the following, we develop a model that implements this concept by an ensemble of oscillators with a learning rule which attunes them to a given sequence. We analyze the error detection accuracy of the model and compare it to those from a cohort of human participants who performed the same sequence learning task. Finally we explore an extension of the model that demonstrates learning of sequences that involve more than one sensory modality.

## 2 Methods

### 2.1 Oscillator ensemble model

We start by developing the model equations for input from a single sensory modality. The learning objective for the ensemble is to associate a set of target inputs *Î* = {*I*_1_, *I*_2_,…} with target phases 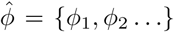. This requires adjusting oscillation frequencies *f* to match the rhythm at which target inputs are presented.

#### 2.1.1 Learning algorithm for tuning individual oscillators

We distinguish three states depending on the phase when an input is presented at time *t* to the oscillator: If the phase *ϕ*(*t*) is close to the target phase 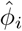 of an input *Î*_*i*_, we call this oscillation *locked* to the rhythm of this input. This is the dynamically stable state for an oscillator, when no further adjustments to its phase or frequency are made by the learning algorithm. If the phase is in a given range around the target phase but not (yet) locked, we call this state *locking*. Oscillations in this state will have their phases set to the target phase of the respective input in the next time step, and the frequency will be adjusted to match the rhythm of the input. We will call any other phase *in transit*, which means that this oscillator will not be tuned in the current time step. Using two corresponding thresholds *θ*_*locked*_ and *θ*_*locking*_, the three states can be formally defined by:

1. locked: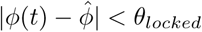
2. locking: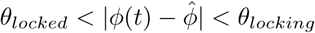
3. in transit: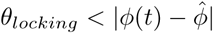

The model runs in discrete time. In each time step, *ϕ* and *f* of every oscillator in the ensemble are updated according to the following equations:

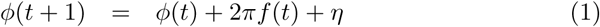

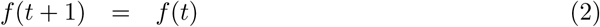

The noise *η* models random fluctuations in the period of neuronal oscillations and is sampled from a normal distribution.

The parameters of oscillators with phases that are locked to the target input or in transit at a given time step are not further modified. Oscillators in the locking state however have their phases and frequencies adjusted depending on the input *I*. If *I* = *Î*_*i*_, the phase is set to the target phase 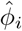 and the frequency is increased or decreased depending on whether the current phase is lagging or leading w.r.t. the target phase:

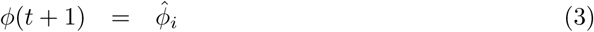

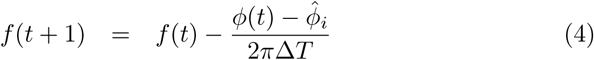

Delta *T* is the number of time steps since the last phase reset of the respective oscillator. It is used to scale the magnitude of the frequency change that is calculated from the phase difference to the magnitude of the oscillator’s current frequency *f* (*t*).

If the input does not correspond to the phase to which an oscillator is locking, i.e., *I* ≠ *Î*_*i*_, then the phase is inverted and the period length is increased or decreased by one time step depending on whether the current phase is lagging or leading w.r.t. the target phase:

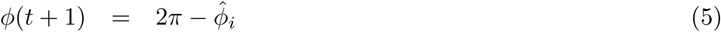

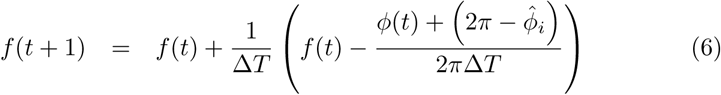

Note that this learning algorithm neither ensures that all rhythms composed by a sequence are picked up by the ensemble nor that the tuning process converges for each oscillator. It does ensure however that the number of locked oscillators monotonically increases over time. The number of rhythms that are picked up from the polyphony in the sequence by the ensemble is a function of the ensemble size, i.e., the number of oscillators.

#### 2.1.2 Calculating the error signal

Initially, most oscillators will adjust their phases and frequencies until they match the rhythm of one of the items in the sequence. As the tuning progresses, fewer and fewer oscillators will be in the *locking* state at any time point. This suggests that the total number of *locking* oscillators is a measure for the attunement of the ensemble to the sequence. Now, if an item suddenly appears at the wrong position, the oscillators that were tuned to the original item at this position would restart tuning, hence the sudden increase in *locking* oscillators could be used to detect incongruent items.

One approach for this detection would be the definition of a threshold which would signal a sequence violation when exceeded. The two problems with this approach are that it is not obvious how such threshold could be defined in advance and that the error signal very likely is above the threshold not only for an incongruent item, but also during the initial learning phase. We therefore looked for a solution that does not require an additional parameter and that accounts for the tuning during the learning phase. What differentiates the learning phase from the re-tuning for an incongruent item is the time since the last phase reset: The initially random phase and frequency of an oscillator will be relatively far off the rhythms that are generated by the sequence; therefore, they will be adjusted several times until they match the rhythm of a particular item. In contrast, the oscillator probably has been attuned for some time before an incongruent item appears. Thus, the time since the last adjustment was made to the oscillator by the learning algorithm is an indicator whether or not this oscillator was in tune with any one rhythm in the sequence. This indicator yields a much stronger signal when an incongruent item perturbs an attuned ensemble than during the initial tuning process. Using the function *δ*_*i*_(*t*) to indicate whether oscillator *i* in an ensemble of size *N* has a phase reset at time *t* (eqs. 3,5), we define the error signal by:

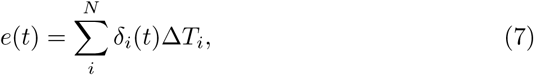

and the decision about the (in-)congruence of the current item is given by

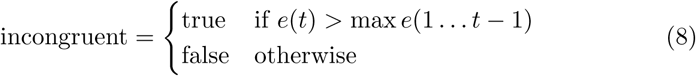

### 2.2 Accommodating several sensory modalities

In the brain, signals from different sensory modalities are processed in different yet interacting cortical areas. We model these cortical areas by modules of oscillator ensembles which receive input from a single modality. Just tagging ensembles as ‘visual’ or ‘auditory’ obviously changes nothing in the dynamics of the corresponding oscillators; therefore, a non-trivial extension of the model towards multimodal sensory input requires introducing additional distinguishing features. Rather than assuming fundamentally different processing mechanisms in different sensory modalities, we consider it to be more appropriate to think of similar mechanisms that operate in different parameter regimes for each modality. For example, auditory processing in the human brain has a higher temporal resolution than visual processing [9], but the anatomical structure of auditory and visual cortices does not seem to be fundamentally different [19]. This finding inspired us to use different base frequencies in different modules. Thus the multimodal model we investigate here consisted of a visual module and an auditory module in which the oscillator ensembles were initialized in a frequency band that was five times higher than that for the ensembles in the visual module. The admittedly arbitrary selection of this frequency ratio was inspired by the intent to demonstrate robustness of the model over a wide range of frequencies.

### 2.3 Numerical simulation

To model the results from the human study, we generated the input from the pixel values of a sequence of images. Each oscillator in an ensemble received input from the same pixel in the images, and there was one ensemble per pixel. Stimulus images from the human study were downsampled to a resolution of 20 × 20 pixels. There was no topographic mapping of the input or any other spatial layout of the ensembles. The two target inputs (*I*_1_ = *black, I*_2_ = *white*) were associated with phases *ϕ*_1_ = *π/*2 and *ϕ*_2_ = 3*/*2*π* respectively. There was also a background color in the images that provided no input (*I* = 0). The distribution for sampling the noise term in eq. 1 had zero mean and a standard deviation of 1 × 10^−10^. The thresholds for defining locked and locking oscillations were *θ*_*locked*_ = *π/*60 and *θ*_*locking*_ = *π/*6.

The properties of both models were determined by running repeatedly numerical simulations with randomized initial conditions. All the results we present below show the average of 100 runs. Initial frequencies for ensembles in the visual module were drawn from a uniform random distribution in the interval [0.01 1], whereas the interval for ensembles in the auditory module was [5 6]. Initial phases in both modules had a uniformly random distribution in the interval [0 2*π*].

### 2.4 Human study

Human participants were studied in two conditions: In one condition, visual and auditory stimuli were presented simultaneously, but subjects were asked to attend to the sequence only in one sensory modality and neglect the other. Therefore we call this condition the unimodal condition. In the other condition, the items of the sequence were presented either as a visual or auditory stimulus, and subjects were requested to attend to an abstract, modality-independent feature of the stimulus and neglect the modality in which the stimulus was presented. We call this condition the crossmodal condition.

The sequences in the unimodal condition were composed of 5 items showing either a horizontally (H) or vertically (V) oriented Gabor patch (10° visual angle, 0.5 cycles per degree), resulting in a total of 32 different sequences. Each stimulus was displayed for 150 ms and followed by 550 ms of a uniform gray background (-). A sine wave tone was presented simultaneously with the image to both ears of the subject. The frequency was either high (2000 Hz) or low (1800 Hz). Its volume was adjusted to 30 dB above the hearing threshold of the subject. The association between pitch of the tone and orientation of the Gabor patch was fixed in all but the last item of the sequence for each subject and randomized across subjects. Figure 1a shows the sequence -V-H-V-V-V as an example.

**Figure 1:**
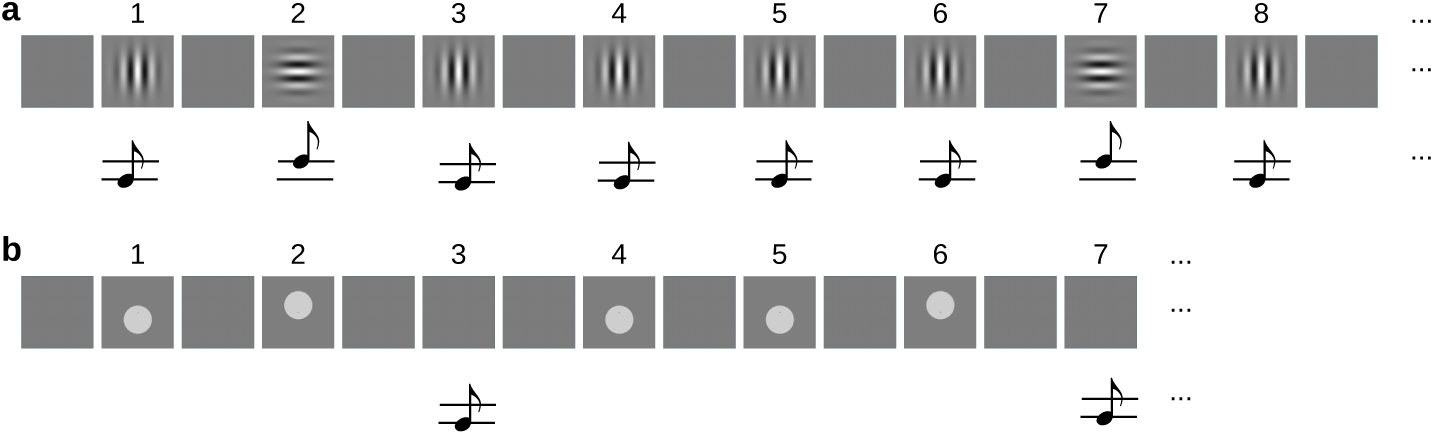
**(a)** Example of an audiovisual sequence for the unimodal task. Sequences were composed of 5 items that were either horizontally (H) or vertically (V) oriented Gabor patches and simultaneously played high- and low-pitch beeps. The sequence -V-H-V-V-V is repeated from item 6 on. **(b)** Example of the sequences that were used in the crossmodal condition. Here, 4 items that were either a visual stimulus (V) or a beep (A) were presented, each representing a ‘high’ (H) or ‘low’ (L) stimulus. The example shows the sequence -VL-VH-AL-VL. The sequence is repeated from item 5 on.

For the crossmodal condition, each item in the sequence was a combination of 2 feature dimensions (height, intensity), 2 feature levels (high/low, strong/weak), and 2 modalities (visual, auditory). Visual ‘high’ and ‘low’ stimuli were gray discs (6° visual angle) above or below the horizontal midline respectively. Auditory stimuli were the same like in the unimodal condition. Intensity was varied between two contrast levels of the disc in the visual stimuli and two volume levels of the beeps. Participants were tested on random subsets from the space of sequences. The trivial sequences in which all items have the same feature level were excluded. In each block of the crossmodal condition, they were requested to attend to only one feature dimension (height or intensity) and neglect the other.

A green fixation cross (0.25° visual angle) was shown at the center of the screen, and subjects were asked to maintain fixation during the stimulation. Sequences were repeated until at least 8 and at most 20 items were presented in the unimodal condition. Within this range, a hazard rate of 0.377 was used to randomize the actual sequence length. Since learning crossmodal sequences was more difficult, at least 10 and at most 20 items were presented in this condition. Here, a hazard rate of 0.448 was used to randomize the actual sequence length. The fixation cross turned red 1200 ms after the offset of the last image, indicating that the subjects should decide whether or not the last item seen was congruent with the sequence. Using the index or middle finger of the right hand, they hit one of two buttons on a response pad that had the responses “yes” (congruent) or “no” (incongruent) assigned. The ratio of congruent/incongruent test items was 0.5. The fixation cross turned green again after the subjects pressed a button, and after another 1500 ms delay, the next trial began.

Sequences were presented in blocks of 32, followed by a short break. Blocks with the congruent/incongruent task were alternated with blocks in which subjects solved an *n*-back memory task. In this task, subjects had to decide whether the last item matched the *n*th previous one. In order to adjust the average performance across participants in the *n*-back memory task to that in the sequence prediction task, 20 of them performed a 1-back task and 9 a 2-back task. In contrast to the congruent/incongruent task, the memory task did not require subjects to learn the whole sequence, but only to remember the last two stimuli seen. In the crossmodal condition, a different control task was employed. Here subjects decided whether or not the last stimulus had appeared anywhere in the sequence before. Deviants were generated by jittering the vertical position of the disc or the pitch of the tone in the terminal stimulus. Each subject completed two sessions of 16 blocks each on separate days.

Twenty nine healthy volunteers (26.3 ± 4.2 years, 17 females) participated in the unimodal human study. Another 25 healthy volunteers (25.1 ± 3.5 years, 14 females) participated in the crossmodal human study. All of them gave written informed consent prior to commencing the experiment. All participants received adequate remuneration. During the experiment, brain activity was recorded by a magnetoencephalograph (MEG). Results of analyzing these neurophysiological data will be published elsewhere. Here we use only the behavioral results to compare them to the properties of the model.

The computational models were studied with the same stimulus material, but the following simplifications were made: The unimodal model was stimulated with the sequence of images only, corresponding to the blocks in which the participants were requested to attend to the visual modality and neglect the auditory. For testing the multimodal model, we used the subset of stimuli that varied only in one feature dimension and that were constant in the other. The model works on a single feature dimension which may be height as well as intensity. Without loss of generality we selected height for the distinguishing feature. From the 256 possible sequences (2 feature levels, 2 modalities, 4 items), we excluded the 32 strictly unimodal ones and tested the model on all remaining 224 truly crossmodal sequences. Figure 1b shows an example sequence.

## 3 Results

### 3.1 Unimodal model

First we demonstrate the properties of the model for two oscillator ensembles which receive input from two representative locations in the images. At location 1 the gray level is different for the horizontal and vertical Gabor patches; at location 2 it is the same (see Fig. 2).

**Figure 2:**
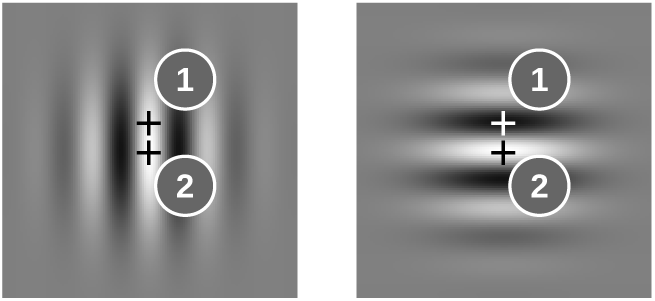
Examples for image locations (marked by ‘+’) where the input to the oscillator ensembles is different for horizontal and vertical Gabor patches (1) and where it is the same (2).

The learning rule adjusts the phases and frequencies to the polyphony that is afforded by the sequence. This attunement process is slower for the more complex input pattern at location 1 than for the regular pattern at location 2, where the frequencies and phases basically converged after about 10 repetitions (Fig. 3a vs. b). The slow attunement in the case of alternating input to the ensemble results from the fact that in the example sequence -H-V-V-V-V, the H stimulus is seen only once per repetition of the sequence (relative frequency of 0.1), and hence more repetitions are needed to synchronize with this input rhythm than to the rhythm of a more frequently presented input. That more oscillators tune to a lower frequency for the input with a less frequent item is reflected in the shallower inclination of the phase plots in Fig. 3c and d.

**Figure 3:**
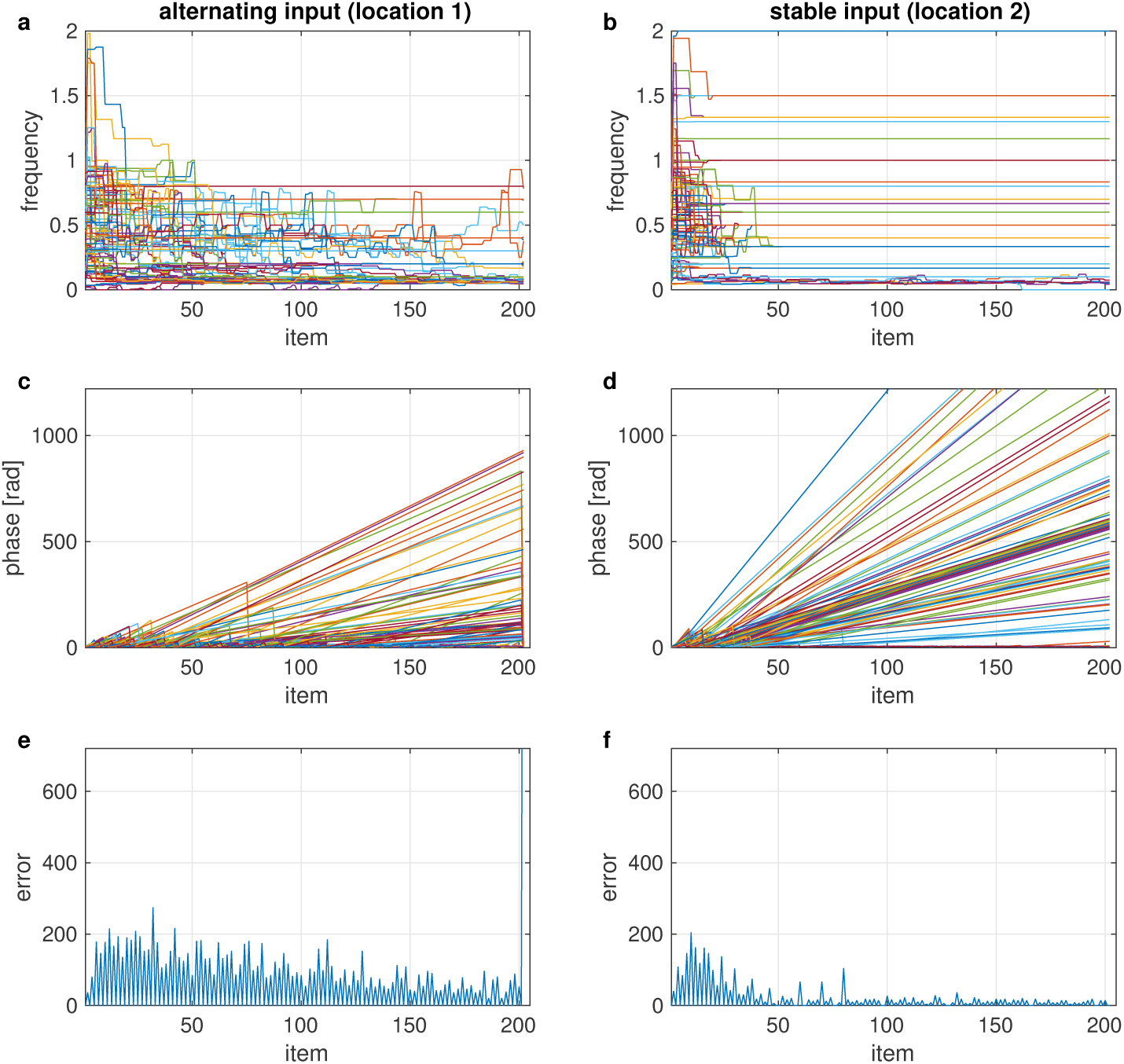
Temporal evolution of frequencies (**a, b**), phases (**c, d**) and error signals (**e, f**) of two ensembles, each consisting of 100 oscillators, with input that differs between items in the sequence (location 1 – **a, c, e**) or is the same in all items (location 2 – **b, d, f**). The sequence was composed of 20 repetitions of -H-V-V-V-V and an incongruent V test stimulus at the end.

If the model is tested with a conflicting item after the sequence was learned, many oscillators in the ensemble undergo a phase reset, which causes a sharp increase of the error signal (Fig. 3e). By detecting whether or not the last item caused a significant increase of the error signal, the model can classify the tested item as incongruent or congruent, respectively.

We analyzed the response accuracy of the model depending on how many times the sequence was repeated before testing an item. Congruence of the tested item is correctly recognized after a few repetitions (Fig. 4a, green curve). Incongruent items, however, seem to require much longer learning time (Fig. 4a, red curve). An interesting observation is that response accuracy for incongruent test items does not increase monotonically with more repetitions, but that it clearly depends on the position of the item in the sequence: It is high when the item at the first position is tested and decreases for the following positions before this pattern is repeated at a higher accuracy level for the next repetition of the sequence. This property is reflected in the periodic modulation of the response accuracy for incongruent items, where the period length is given by the number of items in the sequence.

**Figure 4:**
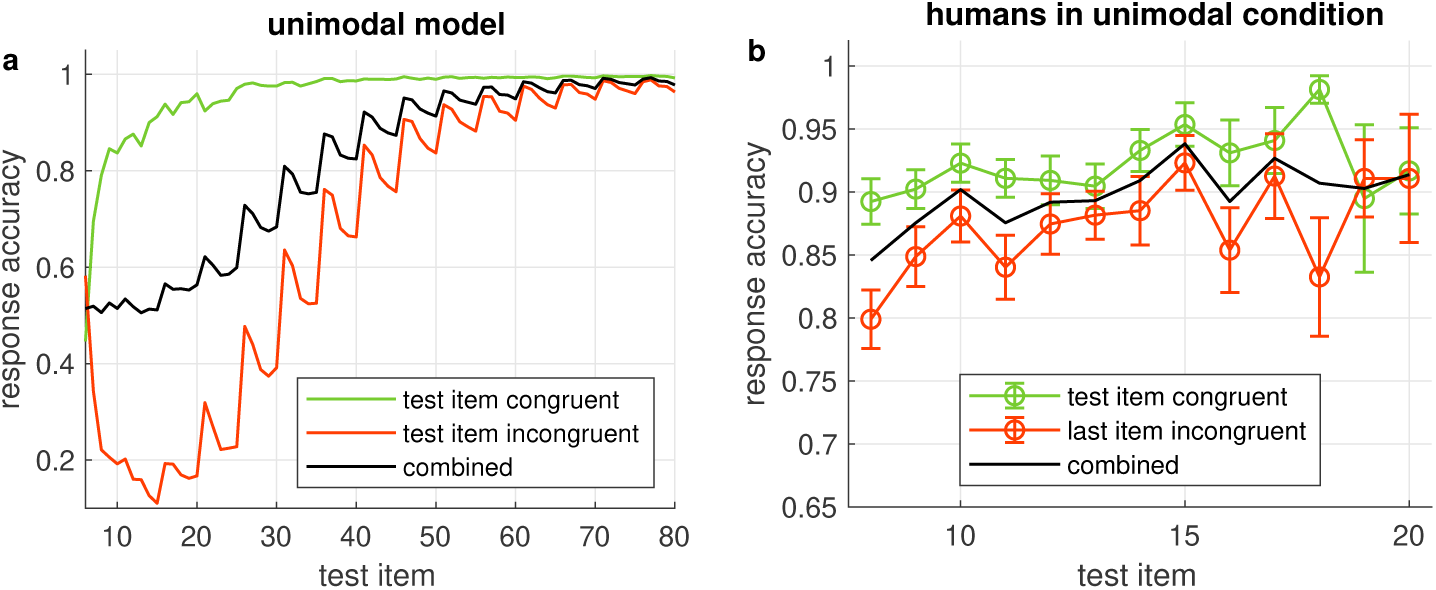
**(a)** Probability of correct model output when the item given on the x-axis is tested in the unimodal model. Green/red curves show the accuracy when the tested item is congruent/incongruent, respectively, black curve is the combined accuracy. All accuracies are averages across all 32 sequences generated from 5 items. **(b)** Average response accuracy of human participants in the unimodal condition. Errorbars show standard error.

After demonstrating the properties of two individual oscillator ensembles, we investigated the dynamic relation between several ensembles. To this end we mapped low-resolution versions of the Gabor stimuli to a corresponding number of oscillator ensembles and analyzed the distribution of the phases and frequencies that developed in the ensembles. Ensembles which received the same input developed similar combinations of phases and frequencies. In Fig. 5 we show the map of phase-frequency clusters that results from the sequence -H-V-V-V-V, for example. After attuning to this sequence, the ensembles developed five clusters with distinct phase-frequency combinations. Clusters of oscillators with the same phase-frequency combination reflect a spatial segmentation of the stimuli in the input sequence.

**Figure 5:**
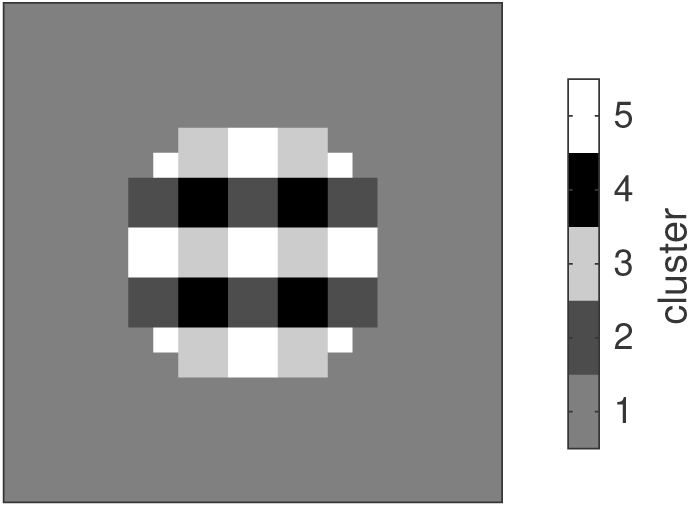
Gray-level-coded map of clusters of oscillators that attune to similar combinations of phase and frequency and hence exhibit a functional coupling.

The distribution of phases and frequencies in each of the five clusters is shown in Fig. 6. Since the image background did not yield any input, the corresponding oscillators retain the initial random distribution of phases and frequencies (cluster 1). Regions with white/black pixels in both stimuli drive the corresponding oscillators to the respective target phases of 3*/*2*π* or *π/*2, respectively (clusters 5 and 4). Most oscillators in these clusters tune to a frequency of 0.5, which reflects the interleaving presentation of an empty stimulus in the sequence. Nevertheless there are oscillators tuning to other frequencies which are compatible with this input rhythm, e.g., 1, 0.3 etc. For image regions where the input alternates between black and white along the sequence, the resulting phase-frequency landscape is more complex. Here the majority of oscillators attune to the frequency of the rare stimulus (H in this example), i.e., 0.1, and the target phase of the respective gray level in this stimulus (cluster 3 - black, cluster 2-white). Other oscillators adjust their phase to the rhythm of the input from the frequent stimulus (V in this example), which is expressed in the phase bins immediately above and below 3*/*2*π* and *π/*2 in clusters 3 and 2, respectively.

**Figure 6:**
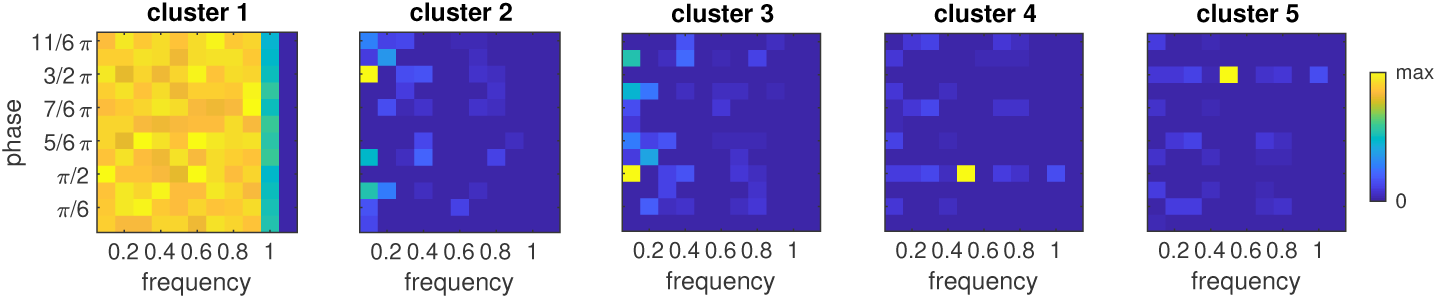
Relative phase-frequency distribution for each of the five clusters shown in Fig. 5. Color represents the number of oscillators with the indicated combination of phase and frequency relative to the bin with the maximum number.

### 3.2 Multimodal model

In a similar manner like for the unimodal model, we investigated the relation between the response accuracy of the multimodal and the number of repetitions of the input sequence. With an increasing number of repetitions, the response accuracy improves, and it is generally higher when congruent items are tested than for incongruent items (Fig. 7a). A comparison of the accuracies with the unimodal model (cf. 4a) shows that the dependence on the sequence repetitions is very similar despite the fact that the multimodal model was tested with a larger variety of sequences (224 vs. 32) which were composed of only four rather than the five elements for the unimodal model.

**Figure 7:**
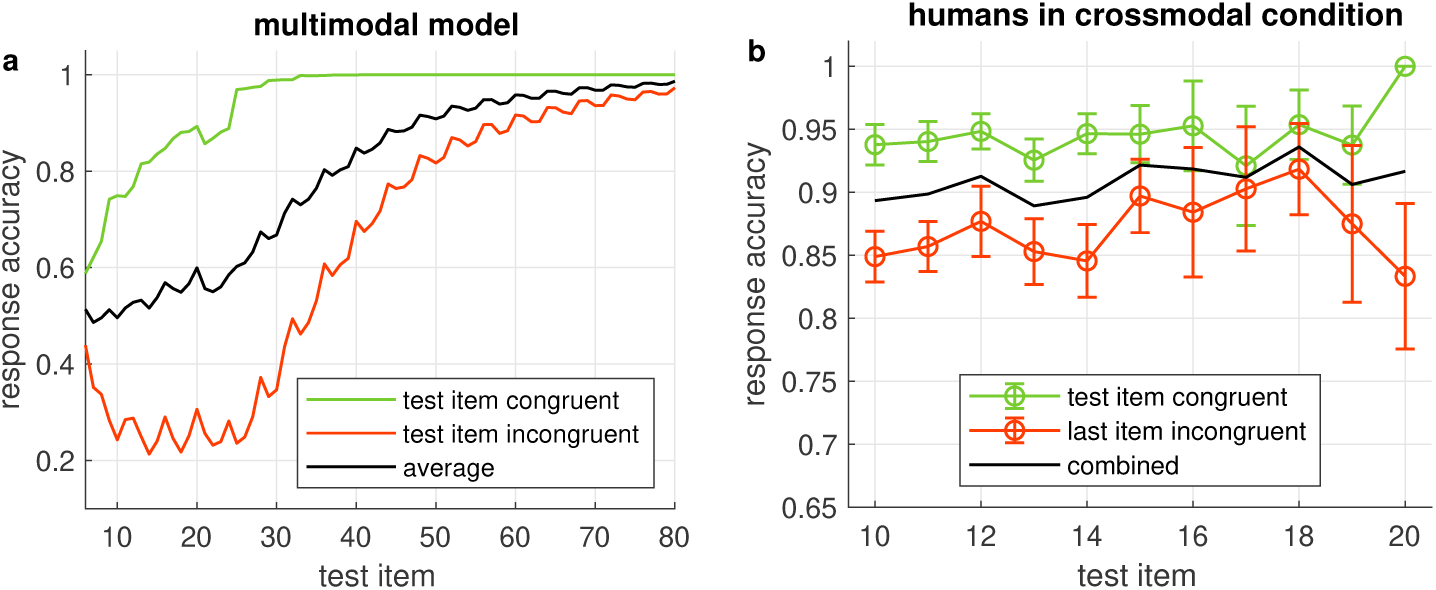
**(a)** Probability of correct model output when the item given on the x-axis is tested in the multimodal model. Green/red curves show the accuracy when the tested item is congruent/incongruent, respectively, black curve is the combined accuracy. All accuracies are averages across all 224 sequences generated from four items. **(b)** Average response accuracy of human participants in the crossmodal condition. Errorbars show standard error.

**Figure 8:**
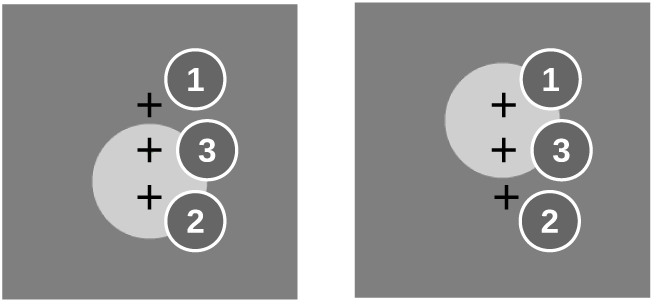
Examples for image locations (marked by ‘+’) in the visual part of the multimodal sequence which provide input only in VH (1), only in VL (2) or both (3) stimuli.

**Figure 9:**
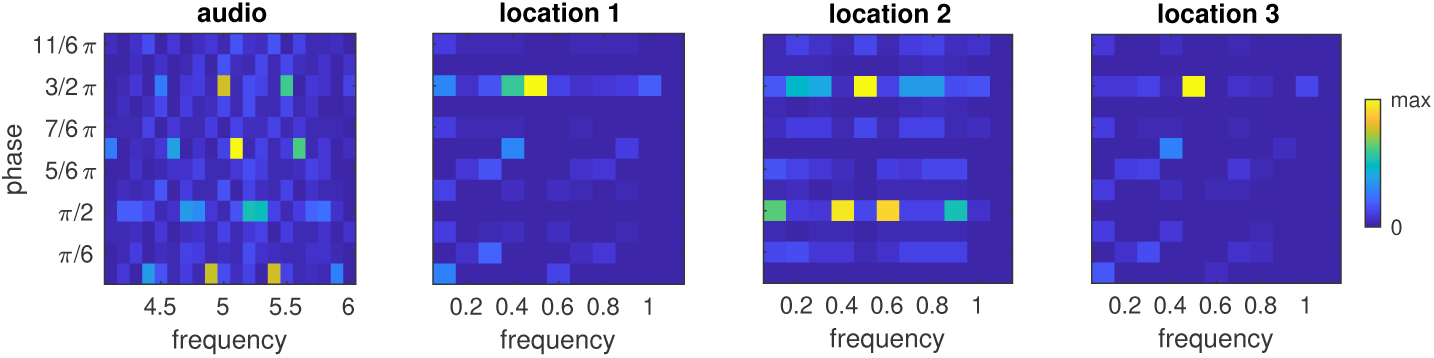
Relative phase-frequency distribution after learning the multimodal sequence AH-VL-VH-VH in an ensemble which received auditory input and three ensembles which received visual input from the representative locations shown in Fig. 8. Color represents the number of oscillators in the cluster that have the indicated combination of phase and frequency. Note the different frequency axes for auditory and visual ensembles.

Finally we considered the distribution of phases and frequencies after a multimodal sequence had been learned. As expected, the majority of oscillators in the ensemble that was stimulated by the auditory signal tuned to the base frequency of the auditory modality (5) and adjusted their phase to the presentation of the auditory stimulus (3*/*2*π*). An interesting finding is that a sizable population of oscillators tuned to the neighboring frequency bins centered around 4.9 and 5.1 and phases of 0 and *π*, respectively. Closer inspection of these phase-frequency combinations revealed that these rhythms never hit the target phase of the auditory stimulus, i.e., they were always in transit when the auditory stimulus appeared, but that their phase nonetheless was compatible with the silent episodes during presentation of the visual stimuli. This pattern of phase-frequency distributions is repeated at the frequencies 4.5 and 5.5.

The ensembles that receive visual input mostly tune to the target phase for bright input (3*/*2*π*) and a frequency of one half the base frequency of the visual modality, i.e., 0.5. In the ensemble that receives input from location 2, several oscillators also tune to the frequencies 0.4 and 0.6 and a phase of *π/*2. This activation of neighboring frequencies at a different phase resembles the observation we made for the auditory ensemble, which likely is a consequence of the fact that the VL stimulus appears at the same frequency in the example sequence as the AH stimulus.

Taken together, the phase-frequency analyses demonstrate that the learning rule tunes the oscillator ensembles to the various rhythms that are generated by repeating the sequence, and that the higher base frequency of the auditory ensemble affords a more complex polyphony to emerge.

### 3.3 Comparison with behavioral results from the human study

Response accuracy of the human participants seemed to increase with more repetitions of the sequence. This trend was more obvious in the unimodal study (Fig. 4b) than in the crossmodal study (Fig. 7b). In both studies, congruent items were more frequently identified correctly than when the tested item was incongruent with the sequence. In comparison with the response accuracies of the models, human performance was always better for a given sequence length and more similar for congruent and incongruent test items. With more sequence repetitions however, the response accuracies of the models increased to the level of the human participants and beyond, indicating that learning is slower in the models.

From the unimodal study, we also analyzed the response accuracies for each of the 32 sequences that the subjects were requested to learn. As expected, the two trivial sequences with only one pattern (always H or V, corresponding to a binary code of 0 and 31, respectively) were the easiest to learn, thus yielding the highest response accuracies (Fig, 10). Next are the sequences in which one element differs from the other four (binary codes 1, 2, 4, 8, 15, 16, 23, 27, 29, 30). The remaining sequences were the most difficult to learn. It is interesting to observe that the response accuracies of the unimodal model largely follow this distribution (Pearson correlation *r* = 0.81, *p* = 2.2 × 10^−8^). The model also reproduces the response accuracies of the human participants when sequences are grouped by complexity quantified by their entropy (Fig, 10, right panel).

**Figure 10:**
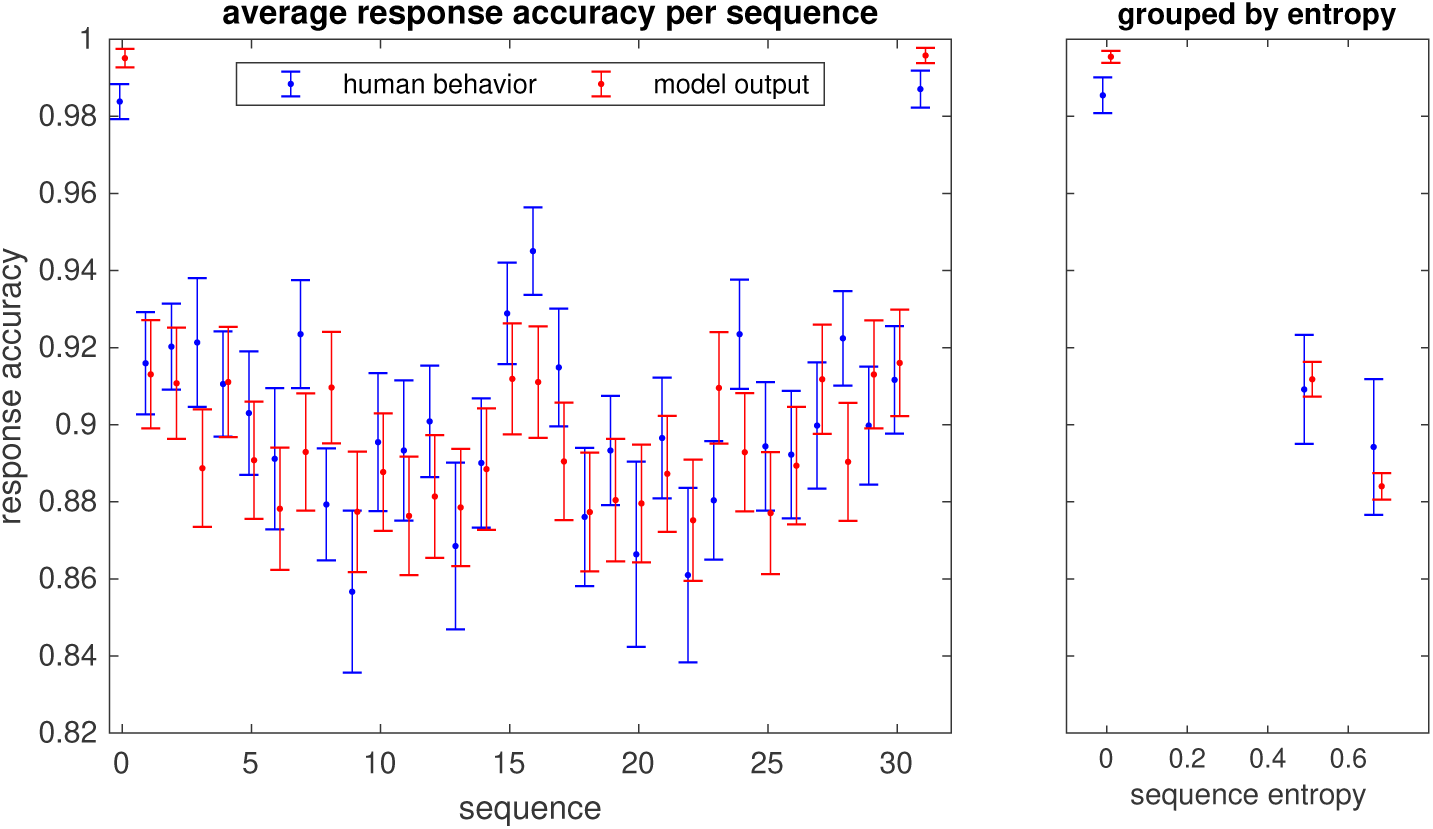
Probabilities of correct response (hit rate) for each of the 32 unimodal sequences that the participants in the study learned (in blue) and response accuracies of the model (in red). Errorbars show the standard error. The sequence number is given by the binary representation of the sequence with the H stimulus corresponding to a 0 bit and V to 1. The right panel shows the average hit rate when sequences are grouped by their entropy.

## 4 Discussion

The oscillator ensemble model is a new approach to sequence learning which exploits the rhythmic, ‘polyphonic’ stimulation that results from repeating a sequence. The basic functional units in this model are oscillators which lock to a rhythm by resetting their phase and adapting their frequency. The results from the unimodal model show that the oscillator ensembles attune to the various rhythms that are generated by a sequence of images. Clusters of distinct combinations of phases and frequencies link image regions that correspond to a meaningful segmentation of the input. Hence clusters of similar phase-frequency distributions can be considered as functional units which link oscillator ensembles that receive input from corresponding regions in visual space. This is an interesting feature, because the segmentation is derived solely from the temporal coherence of image patterns and not from a topographical map of the input. Whereas the functional coupling between ensembles within a cluster is given by their tuning to the same frequency but different phases, such coupling between clusters can be established by oscillators sharing the same phase but having different harmonic frequencies. It has been suggested that such cross-frequency coupling is relevant for integrating functional systems across multiple spatiotemporal scales in the human brain, and it has developed to a well-established concept for understanding brain activity [7]. In our model, cross-frequency coupling is not achieved by fitting the ensemble with a set of fixed frequencies; instead, it results from tuning frequencies and phases to the rhythms in the sequence. The multimodal version of the model demonstrates that the functional coupling also links neuronal populations which operate in different parameter ranges for processing sensory information from different modalities.

It seems also noteworthy that the model does not build or maintain an iconic internal representation of the stimuli. Yet it is capable of predicting whether or not an input is a valid continuation of a sequence. Any incongruent input perturbs the phases of those oscillators that hitherto were attuned to the rhythm of the item at the respective position in the sequence. In the model, this perturbation generates an error signal. The magnitude of this signal is much larger for perturbations of attuned oscillators than for the phase and frequency adjustments made during the initial phase of the learning process. The ability to correctly predict whether or not a given input is a valid continuation of the sequence improves with the number of its repetitions. Our analyses show that the model can correctly identify valid inputs after only a few repetitions, but that the recognition of incongruent inputs requires to repeat the sequence more often. This matches well with the observations from human sequence learning, albeit the models need a longer learning phase to reach the response accuracy of the human participants. Investigating the effect of the model parameters on the learning rate is beyond the scope of the current study. Another aspect that we did not investigate here is that the model could also be used to detect inaccuracies in the timing of the stimulus presentation. It is therefore general enough to cover aspects of predicting ‘what’ and ‘when’ at the same time.

In our model, item position is encoded in the phase relation of a multitude of rhythms which are entrained by the sequence. This corresponds well with concepts for sequence encoding in the hippocampus, derived from animal studies, in which the timing of spikes relative to the phase of ongoing extracellular theta oscillations is considered to encode position in a behavioral sequence. Even if the stimuli are separated by several seconds, their order information is compressed into a single theta cycle, providing a mechanism for short-term buffering and working memory [15]. When the animal traverses a sequence of places, sequence items subsequently move towards the beginning of the theta cycle. This phase precession has been suggested to be the underlying mechanism for episodic memory [14]. In the human brain, the phase relation between gamma and theta oscillations may constitute a similar mechanism [12]. Our model also relates to the multi-timescale, quasi-rhythmic properties of speech, where coordinated delta, theta and gamma oscillations have been suggested to hierarchically structure incoming information [11]. Further support for the relevance of frequency and phase adaptation comes from earlier studies which found single-cell oscillators in somatosensory cortex of awake monkeys that seemed to operate as a phase-locked loop (PLL) for processing of tactile information during texture discrimination [2]. Phase and frequency adaptation has also been observed in thalamo-cortical loops in the brain of rats and guinea pigs, where the frequency of spontaneous oscillations shifted under rhythmic stimulation of a whisker to the stimulation frequency. This may be an essential function for actively decoding information from vibrissal touch [1].

The joint phase space of the oscillators in an ensemble constitutes a pace-maker system that could be used for the discrimination between intervals in the range of seconds, minutes and for circadian rhythm [5]. Even when the oscillation frequencies in the set are in the same range but have slightly different periods, the characteristic ‘beating’, i.e., the time after which the phases of several of these oscillators match, can be exploited to learn sequences of time intervals [17].

By comparing the properties of the model with results of humans in a sequence learning task, we contribute to a long line of approaches to understanding the properties of human sequence learning through the development of oscillator models that reproduce the structure of errors that humans make in sequence learning (see overview in [3, 5]). The main difference between these models and ours is how they explain what drives the oscillator ensemble. Whereas in our model the oscillator rhythms adjust to the sequence, those models work with sets of intrinsically driven, fixed-frequency oscillations. This internal pacemaker provides a dynamic learning context that can be associated with the occurrence of an event by Hebbian learning, for example [3]. It has been argued that models of association with intrinsic oscillation are more compatible with findings from experimental studies on the sequence and timing of events [10]. However, the striking similarity in the structure of errors for congruent and incongruent test items as well as for varying levels of complexity of sequences between the oscillator ensemble model and the human participants in our study suggests that, at least in this dataset, entrained oscillations captured the relevant processes for solving the task. It seems worth therefore to explore the implications of a concept in which externally entrainable and intrinsically driven oscillations interact.

## Author Contributions

AM and AKE conceptualized the research; AM performed experiments and analyzed the data; PW carried out the human study and provided data; AM wrote the manuscript; AKE, JD, and PW revised the manuscript; AKE administrated the project and acquired funding.

## Data Availability Statement

The raw data supporting the conclusions of this manuscript will be made available by the authors, without undue reservation, to any qualified researcher.

